# Adult-Onset Deletion of *ATP13A2* in Mice Induces Progressive Nigrostriatal Pathway Dopaminergic Degeneration and Lysosomal Abnormalities

**DOI:** 10.1101/2024.01.25.577280

**Authors:** Madalynn L. Erb, Kayla Sipple, Nathan Levine, Xi Chen, Darren J. Moore

## Abstract

Although most cases of Parkinson’s disease (PD) are sporadic, mutations in over 20 genes are known to cause heritable forms of PD. A surprising number of familial PD-linked genes and PD risk genes are involved in intracellular trafficking and protein degradation. Recessive loss-of-function mutations in *ATP13A2*, a lysosomal transmembrane P5_B_-type ATPase and polyamine exporter, can cause early-onset familial PD. Familial *ATP13A2* mutations are also linked to related neurodegenerative diseases, including Kufor-Rakeb syndrome (KRS), hereditary spastic paraplegias (HSPs), neuronal ceroid lipofuscinosis, and amyotrophic lateral sclerosis (ALS). Given the severe effects of *ATP13A2* mutations in humans, it is surprising that *ATP13A2* knockout (KO) mice fail to exhibit neurodegeneration even at advanced ages. This discrepancy between human subjects and rodents makes it challenging to study the neuropathological effects of ATP13A2 loss *in vivo*. Germline deletion of *ATP13A2* in rodents may trigger the upregulation of compensatory pathways during embryonic development that mask the full neurotoxic effects of ATP13A2 loss in the brain. To explore this idea, we selectively deleted *ATP13A2* in the adult mouse brain by the unilateral delivery of an AAV-Cre vector into the substantia nigra of young adult mice carrying conditional *lox*P-flanked *ATP13A2* KO alleles. We observe a progressive loss of striatal dopaminergic nerve terminals at 3 and 10 months after AAV-Cre delivery. Cre-injected mice also exhibit robust dopaminergic neuronal degeneration in the substantia nigra at 10 months. Adult-onset *ATP13A2* KO also recreates many of the phenotypes observed in aged germline *ATP13A2* KO mice, including lysosomal abnormalities, p62-positive inclusions, and neuroinflammation. Our study demonstrates that the adult-onset homozygous deletion of *ATP13A2* in the nigrostriatal pathway produces robust and progressive dopaminergic neurodegeneration that serves as a useful *in vivo* model of *ATP13A2*-related neurodegenerative diseases.

## Introduction

Parkinson’s disease (PD) is the most common neurodegenerative movement disorder. PD is characterized by the progressive and irreversible loss of dopaminergic neurons in the substantia nigra pars compacta (SNpc) resulting in a variety of motor symptoms, including resting tremor, rigidity, bradykinesia, and postural instability. Most cases of PD are sporadic and of unknown cause. Risk for developing sporadic PD is thought to result from a combination of genetic and environmental factors together with aging. Interestingly, 5-10% of PD cases are familial and are known to be linked to inherited mutations in single genes i.e. monogenic PD^1,2^.

At this time, mutations in 21 genes have been identified as the cause of familial forms of PD^3^. These genes offer insights into putative molecular disease pathways and provide opportunities to develop animal models that recreate the progressive neuropathology and degeneration in the nigrostriatal pathway that characterizes PD. A surprising number of familial PD genes are involved in intracellular trafficking and protein degradation, implicating these pathways in PD pathogenesis^4,5^. Homozygous loss-of-function mutations in *ATP13A2* cause autosomal recessive, juvenile-onset atypical parkinsonism whereas heterozygous mutations in *ATP13A2* have also been linked to early-onset familial PD^6–8^. Interestingly, ATP13A2 mRNA and protein levels are widely expressed throughout the mammalian brain in multiple cell types and are elevated in the SNpc and other brain regions of sporadic PD subjects^9,10^. *ATP13A2* mutations have also been linked to other familial neurodegenerative diseases, including Kufor-Rakeb syndrome (KRS), hereditary spastic paraplegias (HSPs), amyotrophic lateral sclerosis (ALS), neurodegeneration with brain iron accumulation (NBIA) and neuronal ceroid lipofuscinosis (NCL)^11–16^.

ATP13A2 is a lysosomal transmembrane P5_B_-type ATPase that can function as a polyamine transporter^9,17–20^. ATP13A2 uses ATP hydrolysis to preferentially pump spermine and spermidine from the lumen of late endosomes and lysosomes into the cytosol, thereby regulating polyamine homeostasis in cells^17^. *ATP13A2* disease-linked mutations or depletion of ATP13A2 protein is known to disrupt lysosomal function in cells, causing the accumulation of lysosomes, lysosomal swelling, reduced lysosomal acidity and decreased lysosomal degradative capacity^21,22^. Given the severe and early-onset impact of *ATP13A2* mutations in humans, one would expect to observe robust neurodegeneration in *ATP13A2* knockout (KO) mice. Surprisingly, germline *ATP13A2* KO mice do not exhibit dopaminergic neurodegeneration in the nigrostriatal pathway or noticeable atrophy in other brain regions, even with advanced age^23,24^. Aged germline *ATP13A2* KO mice exhibit modest motor symptoms as well as reactive astrogliosis throughout the brain^23,24^. Additionally, *ATP13A2* KO mice show signs of impaired lysosomal and autophagic function in the brain that become more severe with age, including the accumulation of lysosomal proteins LAMP1 and LAMP2, the accumulation of lipofuscin and ubiquitin aggregation^23,24^.

The discrepancy between human subjects and rodent models bearing homozygous *ATP13A2* mutations or deletions makes it challenging to study the neuropathological effects of ATP13A2 loss *in vivo*. We hypothesize that germline deletion of *ATP13A2* in rodents may trigger the upregulation of compensatory pathways during embryonic development that mask the full neurotoxic effects of *ATP13A2* KO in the brain. Depleting ATP13A2 protein from the mature adult brain, which is likely less resilient and plastic to disruptions in critical molecular pathways, could potentially recreate neurotoxic effects similar to those observed in human subjects. To deplete ATP13A2 selectively in the nigrostriatal pathway of adult mice, we unilaterally delivered AAV-Cre vectors to the SNpc of young adult mice bearing conditional *lox*P-flanked *ATP13A2* KO alleles. Conditional KO mice were assessed at 3 or 10 months after Cre recombinase delivery to monitor the genomic KO of *ATP13A2*, dopaminergic neurodegeneration, axonal degeneration, pathological protein aggregation, neuroinflammation and lysosomal abnormalities.

## Methods

### Animals

Homozygous *ATP13A2 lox*P-flanked (floxed) KO mice (JAX strain # 028387), containing floxed exons 2-3 (**Fig. 1B**), were originally described by Kett *et al.* (2015) and obtained from The Jackson Laboratory^23^. *ATP13A2* floxed KO alleles were genotyped by PCR with genomic DNA^23^. Mice were housed in a pathogen-free barrier facility with a 12-h light/dark cycle. Food and water were provided *ad libitum*. Mice were treated in accordance with the NIH Guidelines for the Care and Use of Laboratory Animals. All animal experiments were approved by the Van Andel Institute Animal Care and Use Committee (IACUC).

**Figure 1.**
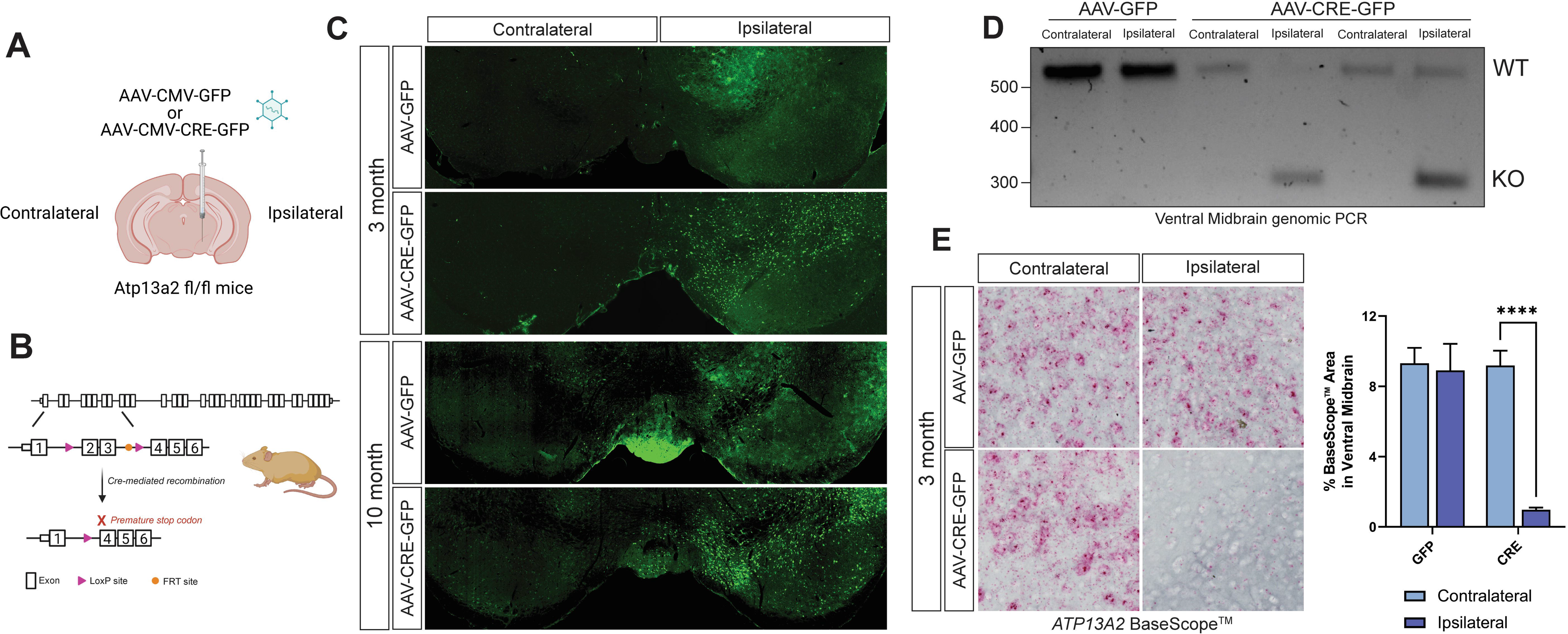
*ATP13A2* knockout in adult mice through unilateral injection of AAV-Cre into the substantia nigra. (**A**) Schematic showing unilateral injection of recombinant AAV-CMV-Cre-GFP or control AAV-CMV-GFP vectors into the substantia nigra in young adult mice. (**B**) Position of *lox*P sites and *FRT* sites in *ATP13A2* floxed KO mice. Cre-mediated recombination results in removal of exons 2 and 3 and formation of a premature stop codon in exon 4. (**C**) GFP or Cre-GFP immunofluorescence in the ipsilateral and contralateral ventral midbrain at 3 or 10 months after AAV delivery. (**D**) Genomic PCR in ventral midbrain tissue of floxed KO mice expressing Cre-GFP or GFP amplifying *ATP13A2* WT and KO alleles using primers flanking exons 2-3. (**E**) BaseScope^TM^ *in situ* hybridization to detect *ATP13A2* mRNA transcript in the ventral midbrain at 3 months. Percent BaseScope-positive area in the ventral midbrain was measured using HALO analysis software. Bars represent mean ± SEM, *n* = 4 animals per group. *\*\*\*\*P*<0.0001 by two-way ANOVA with Sidak’s multiple comparisons test.

### Stereotactic Brain Surgery

Homozygous *ATP13A2* floxed KO mice (age 2-6 months) were anesthetized using 2% isoflurane and positioned in stereotactic frames. We used the following coordinates to administer unilateral injections of AAV vectors into the SNpc: anterior-posterior (A-P), -2.9 mm; medio-lateral (M-L), -1.3 mm; dorso-ventral (D-V), -4.2 mm. AAV vectors (2 µl volume) were delivered at a flow rate of 0.2 µl/min. AAV vectors were purchased from University of North Carolina (UNC) Vector Core. Mice received ∼1.12 × 10^10^ viral genomes (vg) of AAV2/5-CMV-Cre-GFP (Lot# AV4955D) or ∼1.16 × 10^10^ vg of AAV2/5-CMV-GFP (Lot# AV5617B) as a control. Mice were sacrificed at 3, 6 or 10 months after AAV injection. Homozygous ROSA26-LRRK2^R1441C^ mice (JAX strain #026293), containing a floxed STOP cassette upstream of a human R1441C LRRK2 transgene, were unilaterally injected with ∼1.0 × 10^10^ vg of AAV2/5-CMV-Cre-GFP and sacrificed after 12 months for histological analysis.

### BaseScope™ In Situ Hybridization

BaseScope RNA *In Situ* Hybridization was performed following the manufacturer’s instructions (ACD Bio) using a custom probe designed to recognize exons 2-3 of mouse ATP13A2.

### Genomic PCR

Genomic DNA was isolated from ipsilateral and contralateral ventral midbrain hemispheres at 3 months after AAV-GFP or AAV-Cre-GFP injections using the DNeasy Blood and Tissue kit (Qiagen) following the manufacturer’s instructions. Genomic PCR for *ATP13A2* floxed or KO alleles was performed using 100 ng genomic DNA and the Kapa2g Fast HotStart PCR Kit (Roche). PCR primers included a forward primer (5’-CTGCAGCTTCGAGAGGAAAG-3’), one floxed reverse primer (5’-CACTCTGTCCTCAGGCTTTC-3’), and one KO reverse primer (5’-AGGTGGGAATCGGGCTAGAG-3’).

### Immunohistochemistry

Mice were deeply anesthetized followed by transcardial perfusion with 0.9% NaCl and then 4% paraformaldehyde (PFA) in 0.1 M phosphate buffer (PB) at pH 7.4. After perfusion, whole brains were removed and incubated in 4% PFA in 0.1 M PB at 4°C overnight. Brain tissue was cryoprotected using 30% sucrose in 0.1 M PB for ≥ 24 hours before microtome sectioning at 35 µm thickness. Immunohistochemical staining using primary antibodies, biotinylated anti-IgG secondary antibodies (Vector Labs), Vectastain Elite ABC kit (Vector Labs), and 3,3′-diaminobenzidine (DAB; Vector Labs) was performed as described by Dues *et al*. 2023^25^. Midbrain sections immunostained for tyrosine hydroxylase (TH) were incubated in cresyl violet solution for 15 minutes. Primary antibodies used include TH (N300-109; Novus Biological), Iba1 (019-19741; Fujifilm Wako Chemical USA), GFAP (G3893; Millipore Sigma), pS129-α-synuclein (ab51253; Abcam), pSer202/pThr205-Tau (AT8) (MN1020; Thermofisher) and GFP (A-11122; Thermofisher). Secondary antibodies include biotinylated goat-anti rabbit and goat-anti mouse (Vector labs).

For immunofluorescence, brain sections were incubated in primary antibody at 4°C overnight and then in secondary antibody (1:500) conjugated to the appropriate fluorophore for 2 hours. Primary antibodies include TH (N300-109; Novus Biological or ab76442; Abcam), LAMP2 (ab13524; Abcam), p62 (GP62-C; Progen), GFP (A-11122; Thermofisher or 11814460001; Roche or GFP-1010; Aves Labs), GAD67 (MAB5406; Millipore Sigma), parvalbumin (ab11427; Abcam) and TFE3 (ab93808; Abcam). Secondary antibodies were purchased from Thermofisher Scientific. They include: goat-anti rabbit AlexaFluor-488 (A-11008), goat-anti rabbit AlexaFluor-546 (A-11010), goat-anti rabbit AlexaFluor-647 (A-21245), goat-anti mouse AlexaFluor-488 (A-11029), goat-anti mouse AlexaFluor-546 (A-11003), goat-anti rat AlexaFluor-647 (A-21247), goat-anti chicken AlexaFluor-488 (A-11039), and goat-anti chicken AlexaFluor-647 (A-21449).

### Optical density analysis of striatal TH-positive terminals

DAB immunostaining was used to label TH in coronal brain sections containing the striatum. Images were obtained using an Aperio ScanScope XT slide scanner at 20X magnification at a resolution of 0.5 µm/pixel. Mean optical density in the striatum was measured in every 4^th^ section using HALO analysis software (Area quantification module; Indica Labs Inc.). 6-8 sections were analyzed per mouse.

### Stereological quantification of substantia nigra TH-positive neurons

Percent TH-positive neuronal loss and percent total Nissl-positive neuronal loss in the SNpc were estimated using unbiased stereological quantification of TH+ and Nissl+ neurons. For this purpose, we analyzed every 4^th^ serial section of the ventral midbrain using the optical fractionator probe of the StereoInvestigator software (Micro Bright Field Biosciences). Tissue was immunostained for TH and counterstained with 0.1% cresyl violet. Analysis area covered the entire SNpc. Random, systematic sampling was performed using a grid of 120 × 120 µm squares and applying an optical dissector with the dimensions 50 × 50 × 14 µm. During analysis, investigators were blinded to experimental conditions.

### Gallyas silver staining

Gallyas silver staining was performed using the FD NeuroSilver^TM^ Kit II (FD NeuroTechnologies Inc) according to the manufacturer’s instructions.

### Western blot analysis

Striatum or ventral midbrain tissue was homogenized and lysed as described by Mir *et al*. for analysis of phosophorylated Rab proteins^26^. Ice cold RIPA buffer (150 mM NaCl, 50 mM Tris pH 8.5, 1% nonidet P-40, 0.05% sodium deoxycholate, 0.1% SDS) was added to Triton-insoluble pellets and sonicated at 10% amplitude for 15 seconds. Lysates were then centrifuged at 6000 rcf for 10 minutes at 4°C. Protein concentrations were measured in triton-soluble fractions and in RIPA-soluble fractions using a Pierce BCA protein assay following the manufacturer’s instructions (Thermofisher Scientific).

Lysates were mixed with 5X Laemmli sample buffer and incubated at 70°C for 10 minutes. 40-75 µg of protein was resolved on 12.5% or 15% SDS-polyacrylamide gels. Protein was transferred to 0.2 µm nitrocellulose membranes (Amersham) at 20 V overnight. Membranes were blocked in 5% nonfat milk, 20 mM Tris pH 7.5, 150 mM NaCl, 0.1% Tween-20 for 1 h and then incubated in primary antibody in blocking buffer at 4°C overnight. Prior to imaging, membranes were washed and incubated with HRP conjugated secondary antibodies and developed using enhanced chemiluminescence (ECL) or ECL prime (Amersham). Membranes were stripped 2-3 times between primary antibodies, using Restore Westernblot Stripping Buffer (Thermofisher Scientific) according to manufacturer’s instructions. Images were acquired using an Amersham Imager 680 imager and were analyzed using NIH ImageJ (FIJI; v1.53t).

Primary antibodies used for Western blotting include GFP (11814460001; Roche), pThr73-Rab10 (ab230261; Abcam), total Rab10 (8127S; Cell Signaling Technology), pSer106-Rab12 (ab256487; Abcam), total Rab12 (18843-1-AP; Protein Tech), α-synuclein (610787; BD Biosciences), actin (MAB1501, Millipore), LAMP1 (ab24170; Abcam), LAMP2 (ab13524; Abcam), p62 (GP62-C; Progen), cathepsin D (sc-6487-R; Santa Cruz), LC3b (3868; Cell Signaling Technology), ubiquitin (3936; Cell Signaling Technology) and Dynamin-1 (PA1-660; Thermofisher Scientific).

### Statistical analysis

Data was analyzed with GraphPad Prism 9 software by unpaired Student’s *t*-test, paired Student’s *t*-test or two-way ANOVA with Sidak’s multiple comparisons test. Graphs were generated using GraphPad Prism 9 and depict all data as mean ± SEM.

## Results

### Conditional deletion of *ATP13A2* in the substantia nigra of adult mice

To selectively delete *ATP13A2* in the nigrostriatal pathway of adult mice, homozygous *ATP13A2* floxed KO mice (at 2-6 months) were subjected to unilateral stereotactic injection of either recombinant AAV2/5-Cre-GFP (AAV-Cre) vector, or AAV2/5-GFP (AAV-GFP) vector as a control, directly into the SNpc (**Fig. 1A**). Using immunofluorescence for GFP, we find that Cre-GFP or GFP are detectable throughout the ipsilateral ventral midbrain at both 3 and 10 months after AAV injection compared to the contralateral non-injected hemisphere (**Fig. 1C**). Cre-GFP protein is abundant at both timepoints and, as expected, is largely nuclear due to the presence of a nuclear localization signal. GFP protein alone exhibits a more diffuse subcellular localization and is also detected at 3 and 10 months. The GFP fluorescence signal in mice injected with AAV-GFP is moderately less abundant than that in AAV-Cre-GFP mice, which most likely results from its reduced stability due to its increased accessibility for degradation in the cytoplasmic compartment.

*ATP13A2* floxed KO mice contain *lox*P sites flanking exons 2 and 3 (**Fig. 1B**)^23^. Cre-mediated recombination results in the removal of exons 2 and 3 from *ATP13A2* and the introduction of a premature stop codon in exon 4^23^. To confirm the efficiency of genomic recombination at the *ATP13A2* locus in mice, we extracted genomic DNA from ventral midbrain tissues at 3 months after injection of AAV-Cre or AAV-GFP and performed genomic PCR to detect floxed and KO alleles. We detect the *ATP13A2* KO allele exclusively in the ipsilateral ventral midbrain of mice injected with AAV-Cre, compared to the contralateral midbrain (**Fig. 1D**). The floxed allele is detected in both the contralateral and ipsilateral ventral midbrain of mice injected with AAV-GFP as well as in AAV-Cre mice, as expected (**Fig. 1D**). To further confirm successful *ATP13A2* KO, we performed BaseScope^TM^ *in situ* hybridization using a custom probe designed to recognize exons 2-3 of the ATP13A2 mRNA transcript (**Fig. 1E** and **Fig. S1**). At 3 months post-injection with AAV-GFP, mice express ATP13A2 mRNA equivalently throughout the dorsal and ventral midbrain in both ipsilateral and contralateral hemispheres. AAV-Cre-injected mice have similarly high levels of ATP13A2 mRNA in the contralateral ventral midbrain yet exhibit a robust reduction in ATP13A2 mRNA BaseScope^TM^ signal in the ipsilateral ventral midbrain (**Fig. 1E** and **Fig. S1**). Quantitation of BaseScope^TM^ area for ATP13A2 mRNA signal reveals significantly reduced levels of ATP13A2 mRNA only in the ipsilateral ventral midbrain of AAV-Cre-injected mice. Of interest, we observe a marked increase of ATP13A2 mRNA signal in some cells at the periphery of the ipsilateral ventral midbrain region in AAV-Cre-injected mice, but not with AAV-GFP (**Fig. S1**), which we hypothesize may be due to a compensatory upregulation of ATP13A2 in select WT glial cells in response to neuronal-selective *ATP13A2* KO. Together, these data indicate robust and prolonged expression of GFP and Cre-GFP that is largely restricted to the ipsilateral ventral midbrain, with Cre expression inducing efficient genomic recombination and subsequent depletion of ATP13A2 mRNA from the ipsilateral ventral midbrain.

### *ATP13A2* KO induces progressive nigrostriatal pathway dopaminergic neurodegeneration

We next examined dopaminergic neurons in the nigrostriatal pathway at 3 or 10 months after AAV-GFP or AAV-Cre injections. Dopaminergic neurons in the SNpc express tyrosine hydroxylase (TH) and project their axons to medium-sized spiny neurons in the striatum. To first evaluate these axonal projections in the striatum, coronal brain sections were immunostained for TH and optical density was measured comparing the ipsilateral and contralateral striata. After 3 months, we find a significant loss (29.05 ± 6.19%) of TH-positive dopaminergic nerve terminals in the ipsilateral striatum of AAV-Cre-injected mice relative to the contralateral striatum, with no obvious terminal loss (5.27 ± 5.42%) in AAV-GFP-injected mice (**Fig. 2A**). However, in the SNpc at 3 months, we do not find a significant loss of TH-positive dopaminergic (12.67 ± 7.81%) or total Nissl-positive (9.22 ± 7.49%) neurons in mice injected with AAV-Cre (**Fig. 2B**), suggesting that early axonal degeneration in the nigrostriatal pathway occurs prior to obvious nigral dopaminergic neuronal loss.

**Figure 2.**
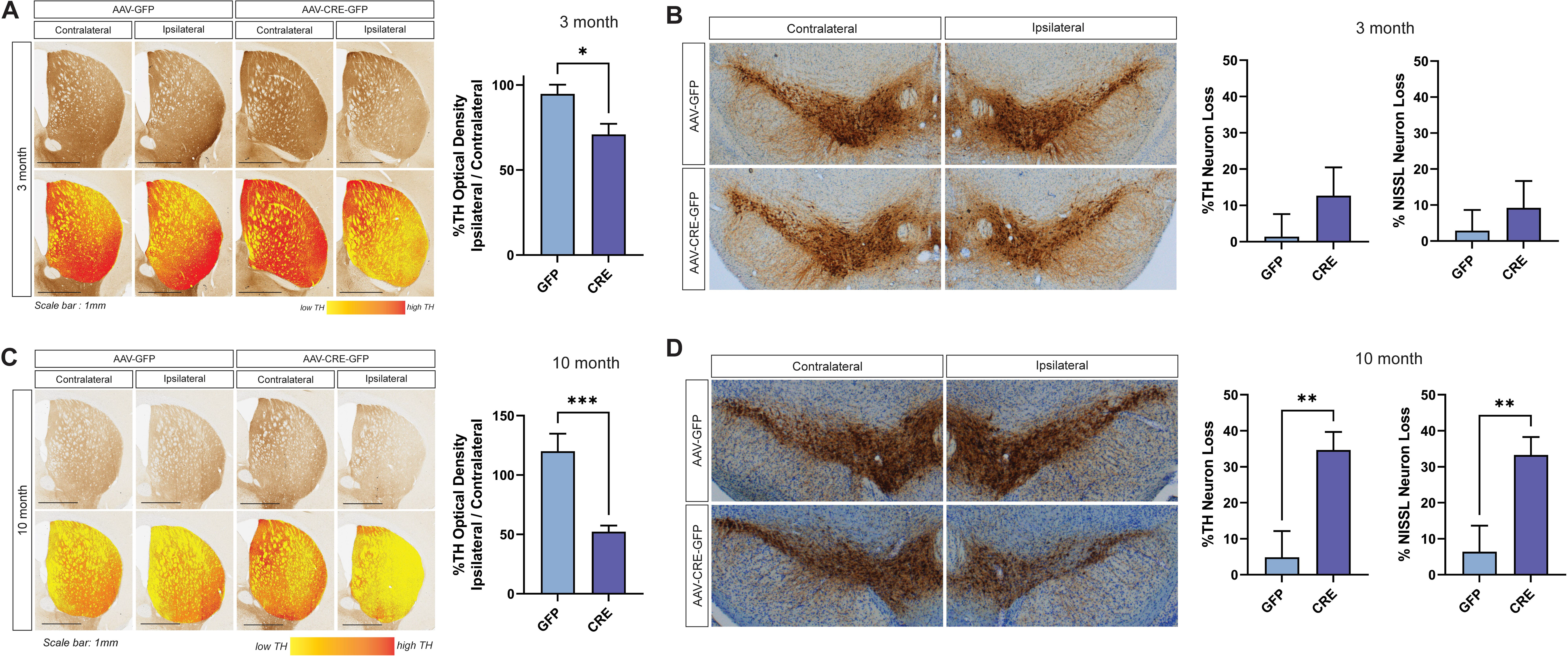
*ATP13A2* knockout causes progressive degeneration of the nigrostriatal dopaminergic pathway over 10 months. (**A**) Representative microscopic images of tyrosine hydroxylase (TH) immunostaining in the striatum of *ATP13A2* floxed KO mice injected with AAV-GFP or AAV-Cre vectors at 3 months. Striatal TH optical density at 3 months was measured using HALO analysis software with data expressed as % TH+ density versus the contralateral hemisphere. Bars represent mean ± SEM, *n* = 7-8 mice per group. *\*P*<0.05 by unpaired, Student’s *t*-test. (**B**) Representative microscopic images of TH immunostaining in the substantia nigra at 3 months. Unbiased stereological analysis of TH+ and Nissl+ neurons in the substantia nigra at 3 months with data expressed as % TH+ or Nissl+ neuron loss versus the contralateral nigra. Bars represent mean ± SEM, *n* = 9-10 mice per group. (**C**) Images of striatal TH immunostaining and striatal TH optical density in mice at 10 months after AAV delivery. Bars represent mean ± SEM, *n* = 13-14 mice per group. *\*\*\*P*<0.001 by unpaired, Student’s *t*-test. (**D**) Images of substantia nigra TH immunostaining and stereological counts of nigral TH+ and Nissl+ neurons at 10 months. Bars represent mean ± SEM, *n* = 12 mice per group. *\*\*P*<0.01 by unpaired, Student’s *t*-test.

At 10 months post-injection, we observe a more robust loss of TH-positive nerve terminals (47.67 ± 5.16%) in the ipsilateral striatum of AAV-Cre-injected mice (**Fig. 2C**), as well as a significant loss of TH-positive dopaminergic (34.68 ± 4.99%) and Nissl-positive neurons (33.29 ± 4.97%) in the ipsilateral SNpc relative to the contralateral hemisphere (**Fig. 2D**). The parallel loss of Nissl-positive neurons confirms neuronal degeneration rather than a loss of TH phenotype. Mice injected with AAV-GFP fail to exhibit any loss of dopaminergic neurons (4.85 ± 7.31%) or their terminals (-20.10 ± 15.93%) at 10 months (**Fig. 2C-D**). These data indicate the progressive degeneration of the nigrostriatal pathway that worsens from 3 to 10 months in the AAV-Cre-injected mice. We also note a qualitative loss of TH-positive dopaminergic neurons (A10 population) in the ipsilateral ventral tegmental area (VTA) of AAV-Cre-injected mice at 10 months (**Fig. 2D**), however, we focused our stereological analysis on SNpc neurons. To determine whether neurodegeneration induced by ATP13A2 depletion is selective to dopaminergic neurons or is non-specific, we immunostained and quantified parvalbumin-positive and GAD67-positive neurons that represent two additional neuronal subpopulations in the substantia nigra (**Fig. S2**). We find no difference in the number of parvalbumin-positive neurons in the ipsilateral ventral midbrain of mice injected with AAV-Cre or AAV-GFP, relative to the contralateral midbrain (**Fig. S2**). Similarly, there is no change in the mean fluorescence intensity of the GAD67-positive neuropil immunostaining in the ipsilateral ventral midbrain of AAV-Cre or AAV-GFP-injected mice (**Fig. S2**). These data suggest that dopaminergic neurons in the ventral midbrain are selectively vulnerable to the neurotoxic effects of ATP13A2 depletion relative to other neuronal subtypes.

One possibility in *ATP13A2* floxed KO mice is that the sustained expression of Cre-GFP in the ventral midbrain over 10 months could independently induce toxicity in nigral dopaminergic neurons. To control for this possibility, we conducted independent experiments by delivering the same AAV-Cre vector into the unilateral SNpc of a different mouse model, ROSA26-LRRK2^R1441C^ conditional transgenic mice, that induces Cre-dependent human R1441C LRRK2 expression (**Fig. S3**). While we observe sustained Cre-GFP expression in the SNpc at 12 months after AAV-Cre delivery, we do not observe any loss of striatal TH-positive nerve terminals or nigral dopaminergic neurons relative to the non-injected contralateral hemisphere (**Fig. S3**). These data indicate that Cre-GFP expression alone is generally well tolerated and is not able to induce neurotoxic effects within the nigrostriatal pathway over these prolonged time periods.

### ATP13A2 depletion induces transient neuroinflammation in the substantia nigra

Germline *ATP13A2* KO mice exhibit reactive astrogliosis as early as 1 month of age, which becomes progressively more severe up to 12-18 months^23^. To examine neuroinflammation in *ATP13A2* floxed KO mice injected with AAV-Cre or AAV-GFP, midbrain sections were immunostained for the astrocyte marker, GFAP, and the microglial marker, Iba1. At 3 months, we find increased GFAP-positive immunolabeling in the ipsilateral ventral midbrain of both AAV-Cre and AAV-GFP-injected mice relative to the contralateral midbrain (**Fig. 3A**). Notably, we observe a significantly larger increase in GFAP signal in the ipsilateral midbrain with AAV-Cre compared to AAV-GFP. At 10 months, there remains a modest yet significant increase in GFAP signal in the ipsilateral ventral midbrain of AAV-Cre-injected mice but no change in AAV-GFP mice (**Fig. 3A**). Increases in GFAP-positive immunolabeling in this KO model appear to result from a combination of AAV-related inflammation and the response to ATP13A2 depletion. Unlike germline *ATP13A2* KO mice, we do not observe progressive astrogliosis as the mice age. Instead, reactive astrogliosis appears to be somewhat transient and is largely attenuated 10 months after AAV-Cre injection.

**Figure 3.**
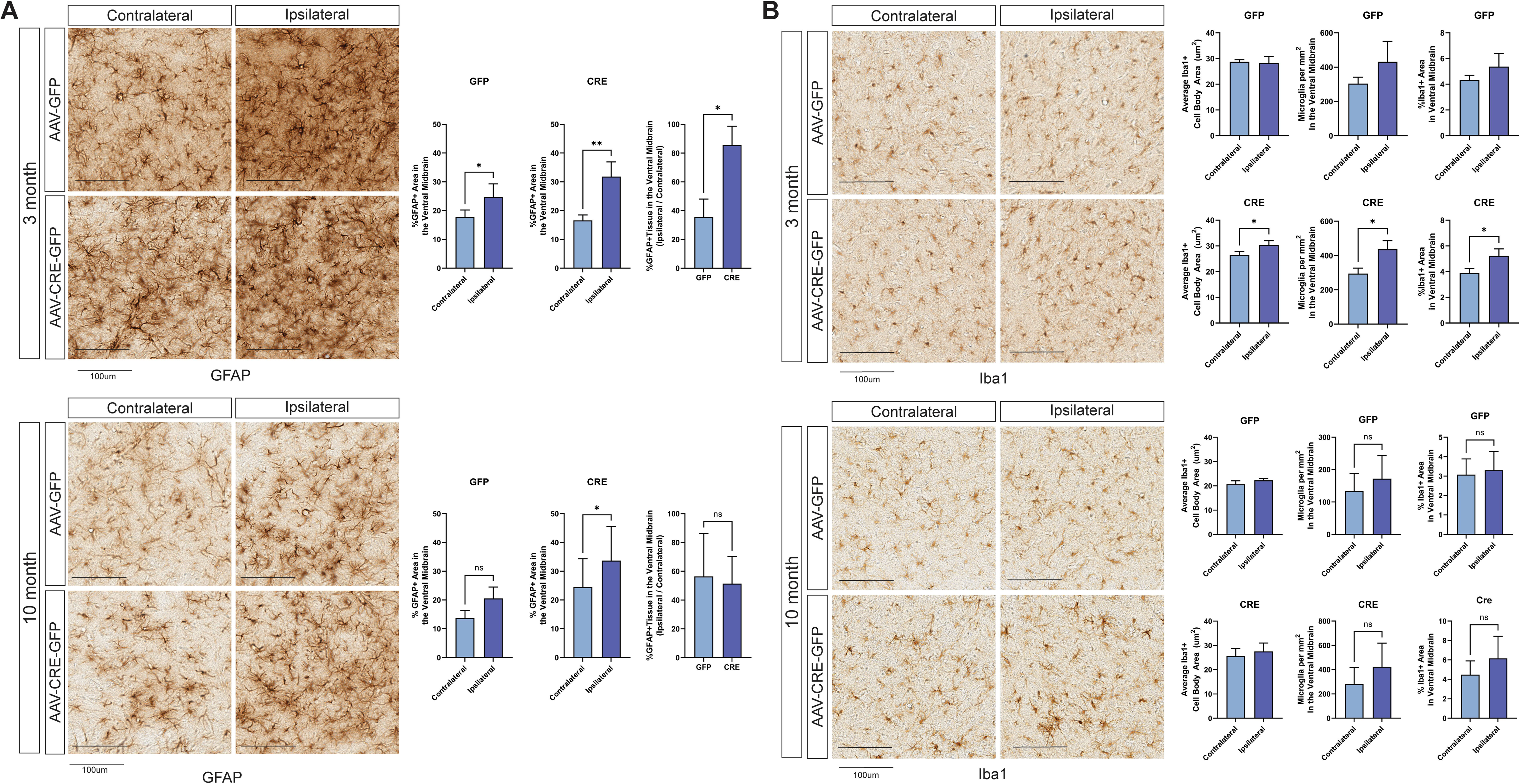
*ATP13A2* KO induces transient neuroinflammation in the ventral midbrain. (**A**) Representative images of GFAP-positive astrocyte immunostaining in the ventral midbrain of *ATP13A2* floxed KO mice unilaterally injected with AAV-GFP or AAV-Cre-GFP vectors at 3 or 10 months. GFAP-positive area was measured in the ipsilateral versus contralateral hemisphere of GFP or Cre-expressing mice using HALO analysis software. Bars represent mean ± SEM, *n* = 8 (at 3 months) or *n* = 4 (at 10 months) mice per group.*\*P*<0.05 or *\*\*P*<0.01 by paired Student’s *t*-test (% GFAP-positive area) or unpaired Student’s *t*-test (% GFAP-positive area, Ipsilateral / Contralateral). (**B**) Representative images of Iba1-positive microglia immunostaining in the ventral midbrain of *ATP13A2* floxed KO mice at 3 or 10 months. Iba1-positive cell number (mm^2^), percent area, or cell body area (µm^2^) was measured in the ipsilateral versus contralateral hemisphere of GFP or Cre-expressing mice using HALO analysis software at 3 or 10 months. Bars represent mean ± SEM, *n* = 4 mice per group. *\*P*<0.05 by paired Student’s *t*-test.

To evaluate microglial activation in these mice, we immunostained midbrain sections for Iba1 and quantified microglial number and morphology (**Fig. 3B**). Microglial activation is characterized by 1) microglial recruitment to the site of neuronal injury, resulting in increased microglial density, and 2) transformation from a resting ramified morphology to an activated ameboid morphology. At 3 months, AAV-GFP-injected mice show no change in Iba1-positive total area, microglial density or microglial cell body area in the ipsilateral ventral midbrain (**Fig. 3B**), suggesting the absence of microglial activation. AAV-Cre-injected mice exhibit a significant increase in Iba1-positive area, microglial density and cell body area, in the ipsilateral ventral midbrain relative to the contralateral hemisphere. Although this microglial activation is relatively modest, it appears to be specific to ATP13A2 depletion, as it does not occur with AAV-GFP injection. Similar to the reactive astrogliosis in these mice, microglial activation is also largely attenuated by 10 months after AAV-Cre delivery (**Fig. 3B**). Therefore, microglial activation also occurs in a transient and early manner.

### Lack of axonal degeneration and protein aggregation in *ATP13A2* KO mice

Given the progressive nigrostriatal dopaminergic pathway degeneration occurring over 10 months in AAV-Cre-injected mice, we evaluated whether axonal damage or degeneration also occurs in these brain regions. Gallyas silver staining was used to label degenerating axons in the striatum or ventral midbrain (**Fig. 4A**). Surprisingly, we do not detect silver-positive degenerating neurites (black fibers) in the ipsilateral striatum or ventral midbrain of AAV-Cre or AAV-GFP-injected mice at 3 or 10 months (**Fig. 4A**). Given the relatively slow progression of TH-positive nerve terminal and cell body degeneration, it is possible that degenerating axons or dendrites are removed from the brain as degeneration occurs, making them challenging to detect.

**Figure 4.**
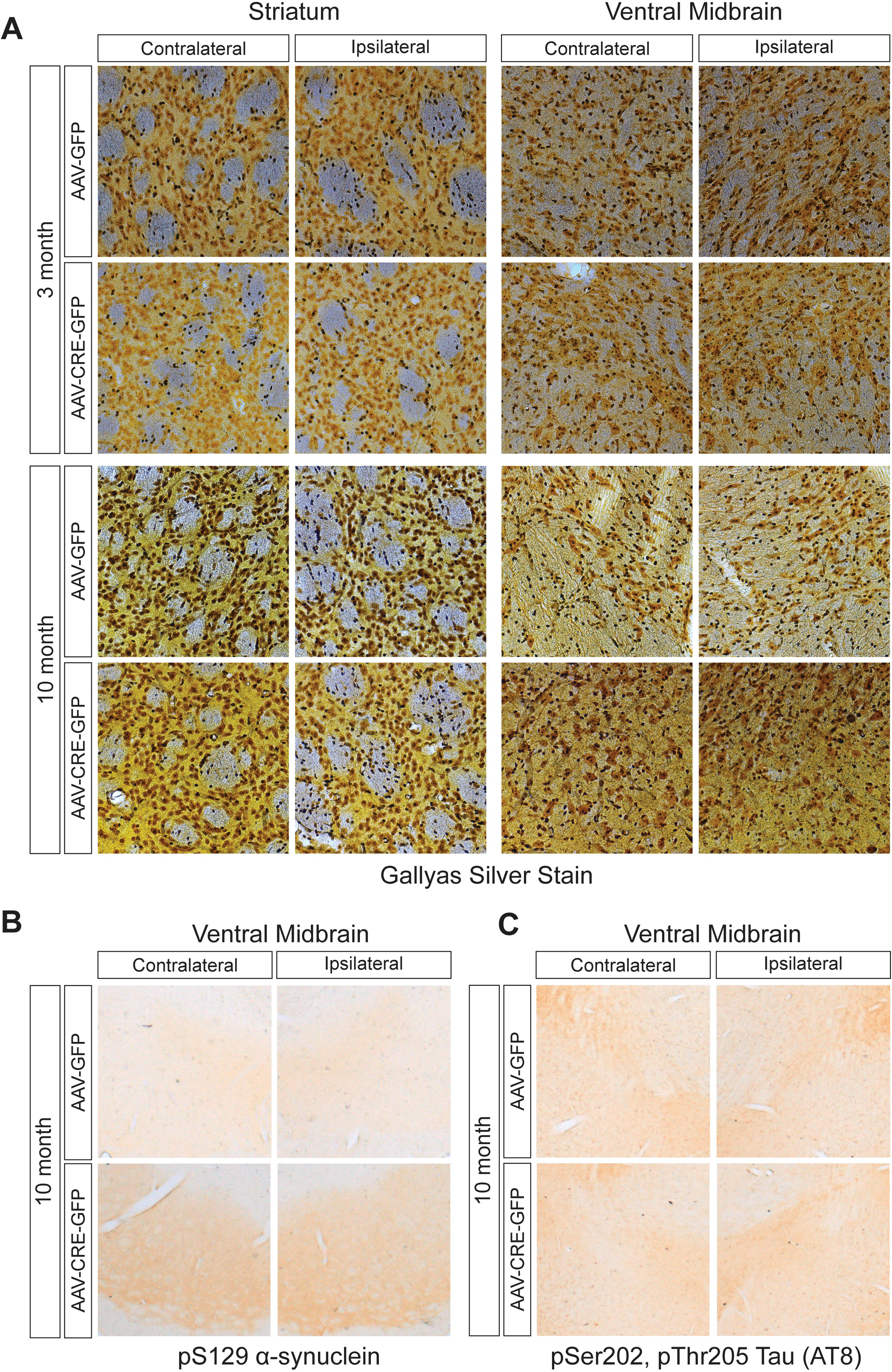
*ATP13A2* KO does not induce axonal damage or accumulation of protein aggregates. (A) Gallyas silver staining was used to detect axonal damage in AAV-Cre-GFP and AAV-GFP-injected *ATP13A2* floxed KO mice. Representative images of striatum and ventral midbrain at 3 or 10 months after AAV injections. (**B**) Ventral midbrain sections from AAV-Cre-GFP or AAV-GFP mice immunostained for pSer129-α-synuclein at 10 months after injections. (**C**) Ventral midbrain sections from AAV-injected mice immunostained for pSer202/pThr205-Tau (AT8) at 10 months after injections.

We next evaluated whether pathological protein aggregation occurs in the conditional *ATP13A2* KO mice. At 10 months, we do not observe the accumulation of pSer129-α-synuclein, a marker of Lewy body pathology, or pSer202/pThr205-tau (AT8), a marker of neurofibrillary tangle pathology, in the ventral midbrain of AAV-Cre-injected mice (**Fig. 4B-C**). These findings are consistent with a previous report that 18-month-old germline *ATP13A2* KO mice do not exhibit increased levels of pSer129-α-synuclein in cortical lysates^23^. Aged germline *ATP13A2* KO mice are reported to exhibit a modest increase in the levels of SDS-soluble α-synuclein in the hippocampus, however, this has not been observed in all *ATP13A2* KO mouse lines and was not observed in the cortex, striatum, midbrain or cerebellum^23,24^. Total tau and huntingtin protein levels are also unchanged in aged germline *ATP13A2* KO mice^24^. At this time, neuropathology has only been reported in one human subject with *ATP13A2*-linked neurodegenerative disease^27^. Interestingly, Chien *et al*. report no aggregation of α-synuclein, pSer202/pThr205-tau (AT8), β-amyloid, TDP43 or p62, in brain tissue from a KRS subject, potentially suggesting that *ATP13A2*-related neuropathology may not involve protein aggregation^27^.

### Lysosomal Abnormalities in *ATP13A2* KO mice

Germline *ATP13A2* KO mice develop pronounced age-dependent lysosomal pathology throughout the brain^23,24^. This lysosomal pathology consists of the accumulation of lysosomal proteins and lipofuscin as well as aggregation of the autophagy proteins ubiquitin and p62/SQSTM1^23,24^. At 10 months after AAV-Cre injection, we observe a significant accumulation of LAMP2-positive lysosomal vesicles specifically in GFP-Cre-positive, TH-positive dopaminergic neurons of the ipsilateral substantia nigra relative to the contralateral nigra (**Fig. 5A**). LAMP2 vesicles also accumulate throughout the ventral midbrain in non-dopaminergic cells. To quantify LAMP2 lysosomal pathology in general, we selectively identified large LAMP2-positive lysosomes, based on fluorescence intensity and lysosome size, in the ipsilateral and contralateral ventral midbrain. We find that ATP13A2 depletion following AAV-Cre delivery leads to a significant accumulation of large LAMP2-positive lysosomes within cells throughout the ipsilateral ventral midbrain (**Fig. 5B**). The number, average size and total area of these enlarged lysosomes are increased in the ipsilateral versus contralateral ventral midbrain (**Fig. 5B**). These data indicate lysosomal abnormalities in dopaminergic neurons and non-dopaminergic cells induced by ATP13A2 depletion throughout the ventral midbrain.

**Figure 5.**
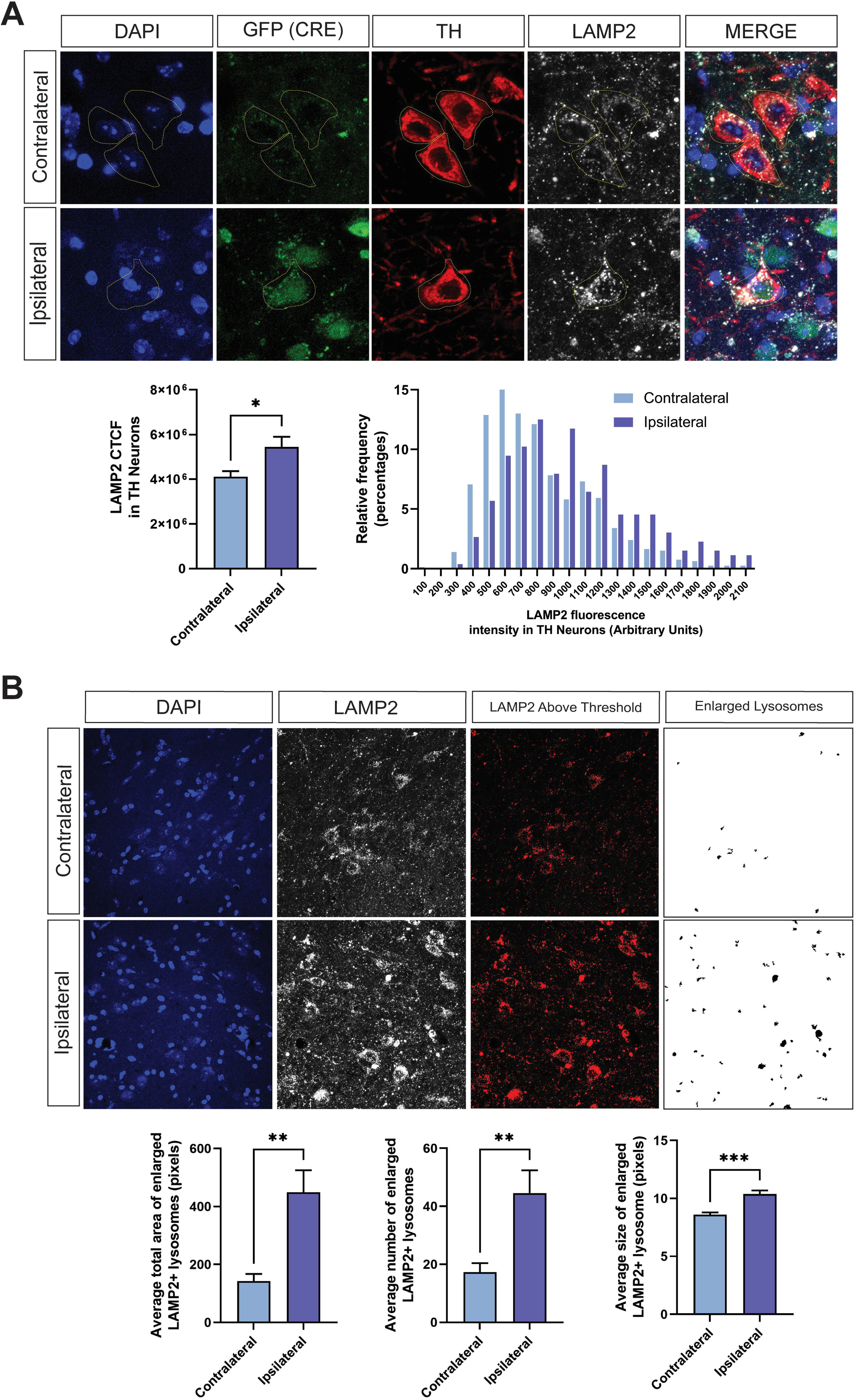
Loss of ATP13A2 leads to the accumulation of LAMP2-positive lysosomes in substantia nigra dopaminergic neurons and lysosomal swelling in the ventral midbrain. (**A**) Confocal immunofluorescent images of LAMP2, TH and GFP (Cre) in ventral midbrain tissue from *ATP13A2* floxed KO mice injected with AAV-Cre-GFP vector after 10 months. LAMP2-positive fluorescence intensity in TH-positive/Cre-GFP-positive cells in the ipsilateral SNpc versus TH-positive cells alone in the contralateral SNpc, was measured using ImageJ analysis software. (*Left graph*) Bars represent mean ± SEM, *n* = 8 mice per group. *\*P*<0.05 by unpaired, Student’s *t*-test. (*Right graph*) Frequency plot indicating the distribution of LAMP2-positive signal intensity in TH-positive neurons of the ipsilateral versus contralateral nigra. Notice the overall rightward shift in frequency in the ipsilateral Cre-GFP-expressing neurons. (**B**) Immunofluorescent images of LAMP2 in ventral midbrain tissue from *ATP13A2* floxed KO mice injected with AAV-Cre-GFP vector after 10 months. The most intense LAMP2-positive signal above an arbitrary threshold within the ventral midbrain region was analyzed using ImageJ software. Intense LAMP2-positive signal was further filtered to remove small LAMP2-positive structures to specifically identify enlarged/swollen lysosomes. The remaining LAMP2-positive lysosomal structures were subjected to a particle analysis to quantify the total area, number, or size of these enlarged lysosomes. Bars represent mean ± SEM, *n* = 8 mice per group. *\*\*P*<0.01 or *\*\*\*P*<0.001 by unpaired, Student’s *t*-test.

Consistent with the phenotype of germline *ATP13A2* KO mice, we find a robust increase in the number of p62-positive inclusions specifically in the ipsilateral ventral midbrain of AAV-Cre-injected mice at 10 months (**Fig. 6A**). p62 is a critical autophagy substrate and adaptor protein that can be used as a reporter to monitor autophagy function, with the formation of p62-positive inclusions indicating autophagy and/or lysosomal disruption^28^. To evaluate lysosomal stress in response to ATP13A2 depletion, we examined the localization of Transcription Factor E3 (TFE3) in the ventral midbrain. Under normal conditions, TFE3 localizes to lysosomal membranes. In response to starvation- or pharmacologically-induced lysosomal stress, TFE3 promotes lysosomal biogenesis by translocating to the nucleus and activating the transcription of a network of lysosomal genes^29^. At 3 months after AAV-Cre injection, we observe strong nuclear localization of TFE3 in a subset of GFP-Cre-positive cells in the ipsilateral SNpc and VTA (**Fig. 6B**). Immunostaining for TH indicates strong TFE3 nuclear localization specifically in TH-positive dopaminergic neurons as well as in surrounding TH-negative cells (**Fig. 6B**). TFE3 nuclear localization is not observed in the contralateral ventral midbrain of AAV-Cre-injected mice, nor in either hemisphere of AAV-GFP-injected mice. Based on the high efficiency of *ATP13A2* knockout throughout the ventral midbrain induced by AAV-Cre, we anticipated more widespread lysosomal stress in this region at 3 months. Surprisingly, we only observe strong TFE3 nuclear localization in a small subset of cells. The timing of the TFE3 response to lysosomal stress in neurons *in vivo* has not been well characterized. Nuclear TFE3 could be a transient response to lysosomal stress in response to ATP13A2 depletion. Alternatively, ATP13A2 depletion may cause a relatively mild form of lysosomal stress that does not consistently activate TFE3. Collectively, these data indicate that a subset of cells in the ventral midbrain, including dopaminergic neurons, exhibit lysosomal activity deficits (p62 accumulation) and stress (nuclear TFE3) induced by the loss of ATP13A2.

**Figure 6.**
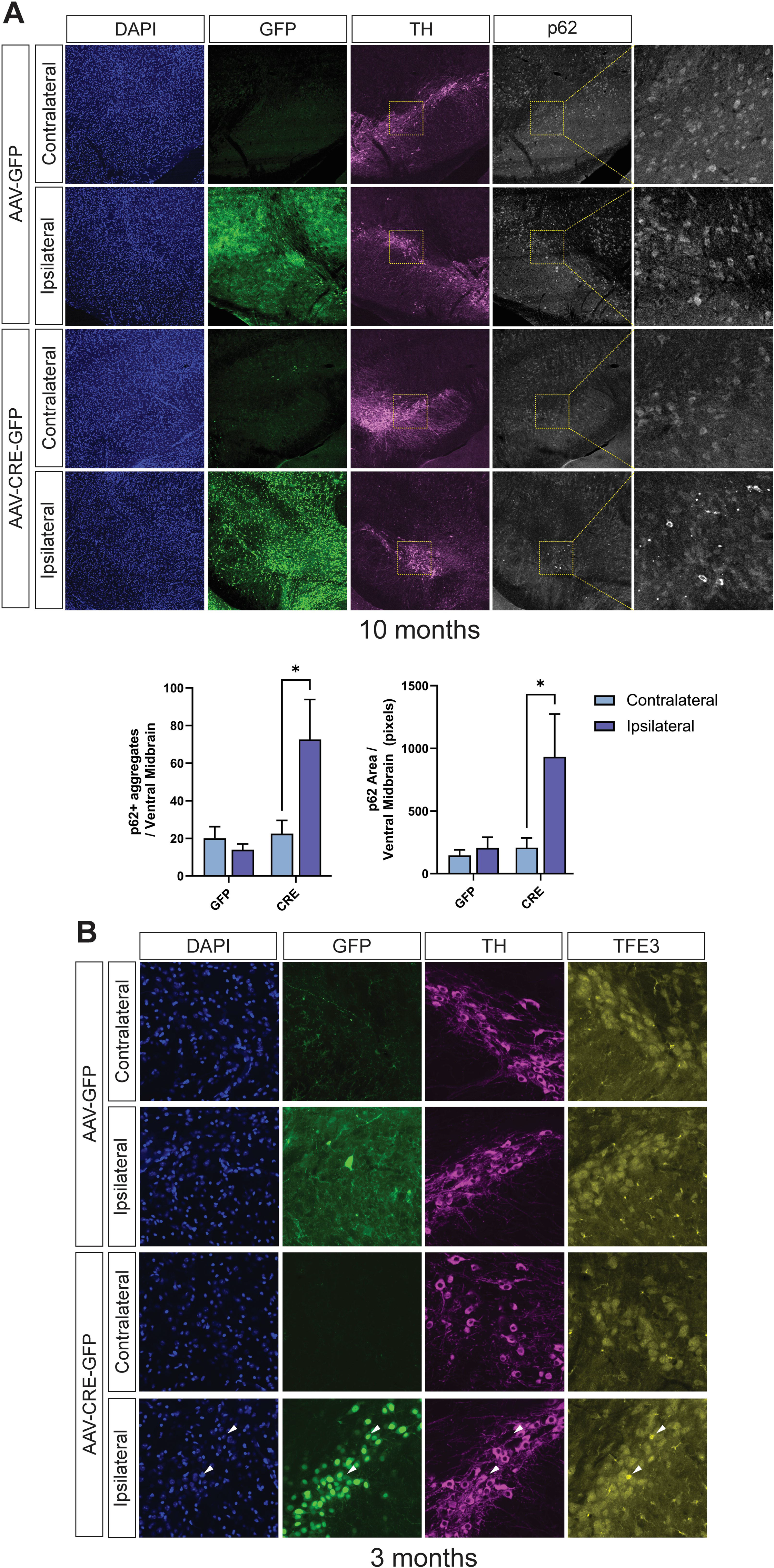
*ATP13A2* deletion disrupts autophagy and induces lysosomal stress. (**A**) Confocal immunofluorescent images of p62, TH and GFP in ventral midbrain tissue from *ATP13A2* floxed KO mice injected with AAV-Cre-GFP or AAV-GFP vectors after 10 months. p62-positive inclusion number and area in the ventral midbrain were measured using Cell Profiler image analysis software. Bars represent mean ± SEM, *n* = 4 mice per group. *\*P*<0.05 by two-way ANOVA with Sidak’s multiple comparisons test. (**B**) Confocal immunofluorescent images of TFE3, TH and GFP in ventral midbrain tissue from *ATP13A2* floxed KO mice injected with AAV-Cre-GFP or AAV-GFP vectors after 3 months. Nuclear translocation of TFE3 (*arrowheads*) was only observed in the ipsilateral ventral midbrain of AAV-Cre-GFP-injected mice. Representative images from *n* = 4 mice per group.

To further evaluate changes in the autophagy-lysosome pathway induced by ATP13A2 loss, we quantified the steady-state levels of key lysosomal and autophagy proteins in soluble ventral midbrain extracts from mice injected with AAV-GFP or AAV-Cre after 3 or 6 months. Surprisingly, we do not find significant changes in the levels of lysosomal or autophagy proteins, including LAMP1, LAMP2, p62, cathepsin D species, LC3B-I and LC3B-II, ubiquitin, phospho-Thr73-Rab10 or phospho-Ser106-Rab12, in the ipsilateral ventral midbrain of AAV-Cre-injected mice (**Fig. S4** and **Fig. S5**). Rab10 and Rab12 are substrates of LRRK2 that can be phosphorylated in response to lysosomal stress or damage^30^. We also do not observe altered levels of total α-synuclein protein in these extracts, that can often accumulate with lysosomal impairment^31^. At 10 months following AAV-Cre injection, we do observe increased LAMP2 signal and p62 aggregation in numerous cells within the ipsilateral ventral midbrain by confocal immunofluorescence (**Fig. 5** and **Fig. 6A**). We suspect that 3 or 6 months after AAV-Cre injection is too early to detect bulk changes in lysosomal proteins in ventral midbrain extracts by Western blotting. Immunofluorescence would likely be more sensitive for detecting subtle changes in lysosomal proteins in individual cells at these timepoints, as we find at 10 months.

## Discussion

The adult-onset KO of *ATP13A2* selectively in the ventral midbrain of mice induced by AAV-Cre delivery replicates many of the phenotypes initially observed in germline *ATP13A2* KO mice, including neuroinflammation and abnormalities in the endolysosomal pathway. Importantly, this approach additionally induces robust and progressive nigrostriatal pathway dopaminergic degeneration over a 10-month period, thereby recapitulating a major neuropathological hallmark of PD. Consistent with germline KO mice, however, the adult-onset deletion of *ATP13A2* does not induce the pathological aggregation of α-synuclein or tau. This is the first *in vivo* mammalian model of ATP13A2 depletion that manifests measurable neurodegeneration. The development of animal models that recreate the molecular pathology and progressive dopaminergic neurodegeneration that characterize PD is one of many critical steps in identifying and evaluating novel disease-modifying therapeutics. This new PD model specifically targets one component of the endolysosomal pathway resulting in a relatively pure and sustained lysosomal dysfunction *in vivo*, that induces neuroinflammation, autophagy dysfunction and eventual dopaminergic neuronal death. Although mutations in *ATP13A2* are relatively rare in human PD subjects, lysosomal dysfunction is observed or implicated in many forms of familial and sporadic PD. Accordingly, this adult-onset *ATP13A2* KO model may be useful in understanding both familial and sporadic forms of the disease, and provides a useful tool for evaluating putative therapeutic strategies targeting the endolysosomal pathway.

The development of dopaminergic neurodegeneration in this adult-onset, conditional *ATP13A2* KO model (**Fig. 2**) contrasts with the lack of neuronal loss in aged germline KO mice^24,28^. We suspect that germline KO mice may upregulate compensatory neuroprotective pathways that preserve dopaminergic neuronal viability, a mechanism that is likely not activated in the adult brain upon conditional *ATP13A2* deletion. A similar phenomenon is observed in *parkin* KO mice, where the Cre-mediated, adult-onset KO of *parkin* is sufficient to induce progressive neurodegeneration whereas germline KO mice consistently fail to do so with advanced age^32–34^. One study has further shown that the lack of neurodegeneration in germline *parkin* KO mice may relate to the upregulation of the mitochondrial pro-survival factor, Mcl-1, and reducing *Mcl-1* gene dosage is sufficient to sensitize *parkin* KO mice to dopaminergic neuronal loss and motor deficits^35,36^. Similar compensatory pathways may be upregulated in germline *ATP13A2* KO mice that mask neurodegeneration, such as lysosomal stress or damage genes, and this would be important to explore in future studies. Newer techniques such as single-nucleus RNA-sequencing of ventral midbrain tissue would permit a comprehensive analysis of both models and provide insight into neuronal susceptibility and specific adaptations to ATP13A2 depletion.

In the *ATP13A2* floxed KO mice, we do observe the nuclear translocation of TFE3 in a small subset of cells in the ventral midbrain (**Fig. 6B**), and this may be sufficient to preserve viability in these cells. It is possible that dopaminergic neurons that eventually degenerate in this KO model are not able to mount an effective response to lysosomal damage by activating the <u>C</u>oordinated <u>L</u>ysosomal <u>E</u>xpression <u>a</u>nd <u>R</u>egulation (CLEAR) gene network via the transcription factors TFE3 or TFEB^37^. The lack of evidence for α-synuclein and tau aggregation in the conditional *ATP13A2* KO mice (**Fig. 4B-C**) is consistent with human neuropathology data from a single *ATP13A2*-linked KRS patient^27^ as well as germline *ATP13A2* KO mice^23^. More importantly, the development of key phenotypes in germline KO mice, such as reactive gliosis, lipofuscinosis, ubiquitinated protein aggregates, and endolysosomal abnormalities, were shown to occur even in the absence of α-synuclein^23^. A similar study in a rat viral-based model of PD revealed that ATP13A2 overexpression was unable to protect against dopaminergic neuronal loss and motor deficits induced by the expression of human wild-type α-synuclein^10^. These data suggest that α-synuclein aggregation may not be a key part of the disease spectrum induced by loss-of-function *ATP13A2* mutations.

The recent characterization of ATP13A2 as a lysosomal polyamine exporter protein suggests that adult-onset *ATP13A2* KO may disrupt polyamine homeostasis in the ventral midbrain^17^. Measuring polyamine levels in ventral midbrain tissue when ATP13A2 is depleted will provide an important confirmation of this activity and insight into this pathway. Additionally, altering polyamine levels in the brain may be sufficient to modulate the pathogenic effects of adult-onset *ATP13A2* KO in nigral dopaminergic neurons. For example, treatment with difluoromethylornithine (DFMO) to inhibit ornithine decarboxylase 1 (ODC1), the rate-limiting enzyme required for polyamine synthesis, would systemically reduce putrescine and spermidine levels in mice^38,39^. Increasing polyamine levels in the brain is more challenging. Dietary supplementation of polyamines can increase circulating polyamine levels but polyamines are not known to efficiently cross the blood brain barrier^40,41^. Since high levels of polyamines cause toxicity in primary neuronal cultures, injecting concentrated polyamines directly into the brain would likely induce toxicity in mice, and may not be a viable experimental strategy^17^. Future studies investigating the effects of altering polyamine levels in the brain of adult-onset *ATP13A2* KO mice could illuminate important aspects of the role of ATP13A2 in polyamine homeostasis *in vivo* and the effects of modulating polyamine levels on endolysosomal pathway-mediated neurotoxicity.

## Availability of Supporting Data

All data generated and analyzed in this study has been presented in this manuscript.

## Author Contributions

MLE designed, performed, and analyzed most experiments. KS assisted with immunohistochemistry and immunofluorescence experiments. NL performed stereotactic surgery, immunohistochemistry and analyzed experiments using ROSA26-LRRK2^R1441C^ mice. NL and XC assisted with stereotactic surgeries. DJM conceptualized and supervised this research. MLE and DJM wrote the manuscript.

## Acknowledgements

We would like to thank the outstanding core facilities at the Van Andel Institute that supported this work, including the Vivarium Core (RRID:SCR 023211), the Pathology and Biorepository Core (RRID:SCR 022912) and the Optical Imaging Core (RRID:SCR 021968). Figure 1A and 1B were made using BioRender.

