## Supplemental Data for "Adult-Onset Deletion of *ATP13A2* in Mice Induces Progressive Nigrostriatal Pathway Dopaminergic Degeneration and Lysosomal Abnormalities"

### Supplemental Figures

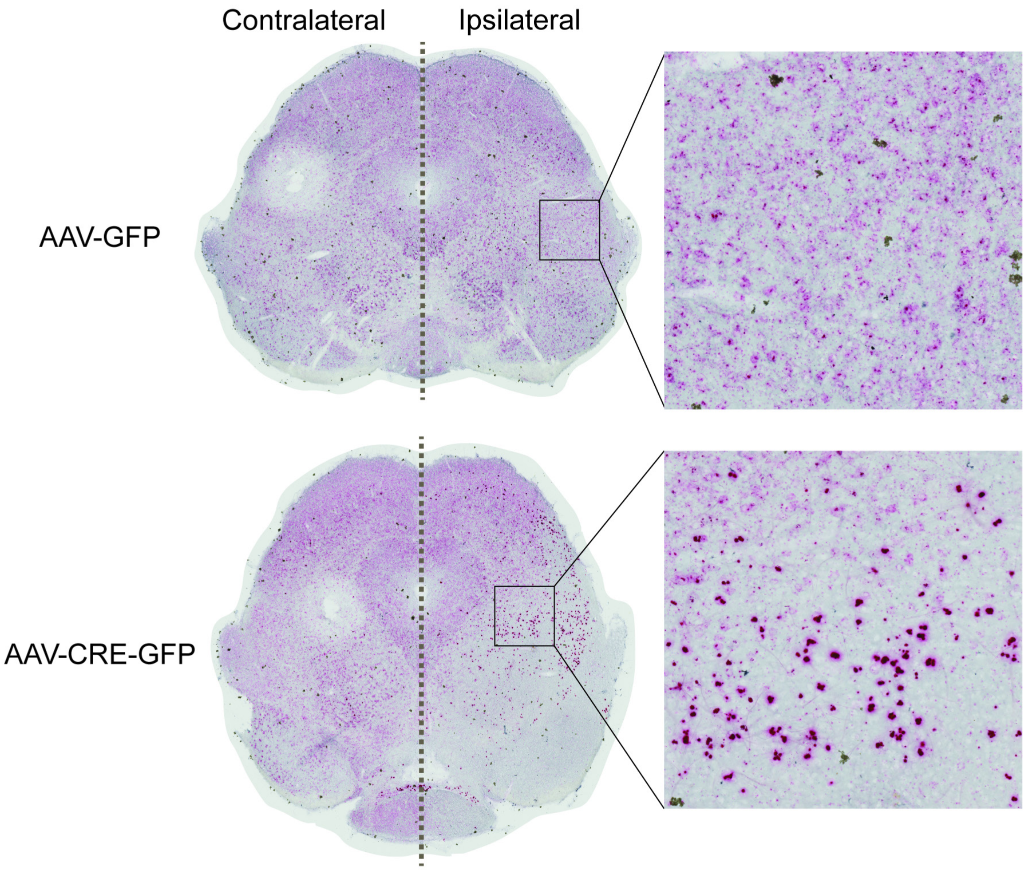


**Figure S1. *ATP13A2* KO in the ventral midbrain induces upregulation of ATP13A2 mRNA transcript in cells at the periphery.** Microscopic colorimetric images of BaseScope™ *in situ* hybridization signal recognizing exons 2-3 of ATP13A2 mRNA transcript in ipsilateral and contralateral midbrain tissue sections of *ATP13A2* floxed KO mice injected with AAV-GFP ($n$ = 3) or AAV-Cre-GFP ($n$ = 4) at 3 months. High power images indicate individual cells from the ipsilateral midbrain. Notice the selective removal of ATP13A2 signal in the ipsilateral ventral midbrain of AAV-Cre-GFP mice, yet a robust increase in signal in some cells at the periphery of the ventral midbrain of these mice.

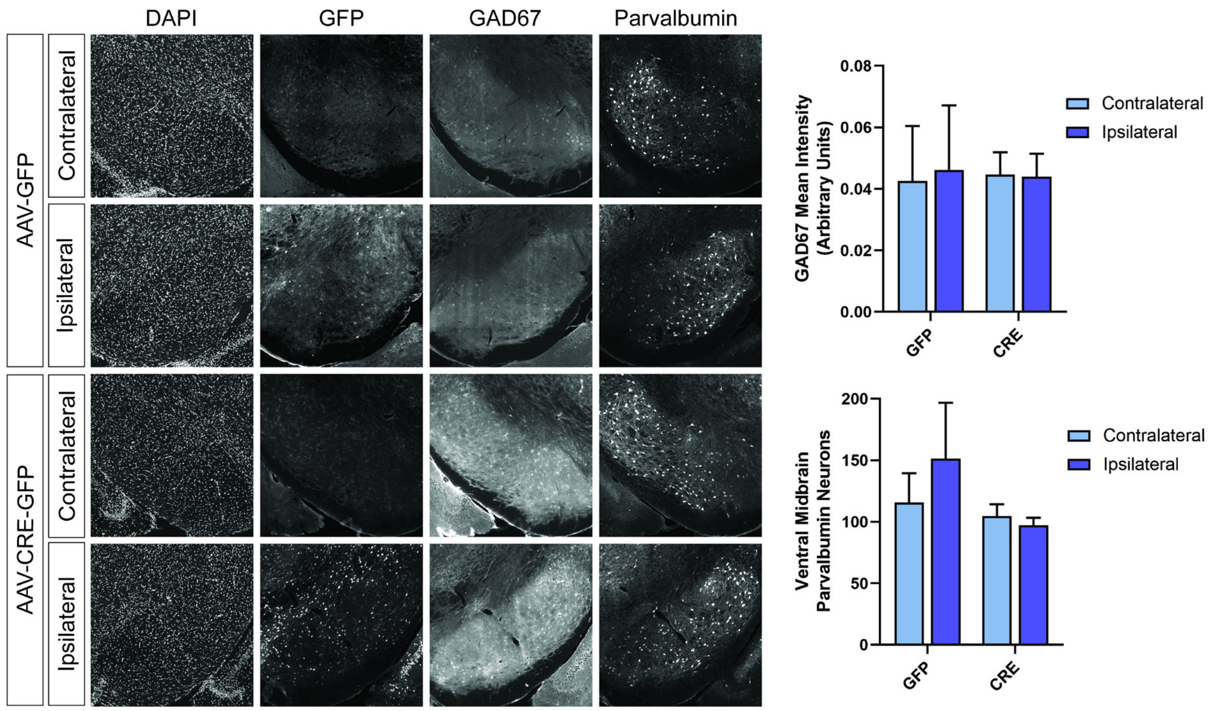


**Figure S2. Loss of *ATP13A2* does not induce degeneration of GAD67-positive or parvalbumin-positive non-dopaminergic neurons in the substantia nigra.** Confocal immunofluorescent images of GAD67, parvalbumin and GFP in the ipsilateral and contralateral substantia nigra from *ATP13A2* floxed KO mice injected with AAV-Cre-GFP or AAV-GFP vectors after 10 months. GAD67-positive neuropil fluorescence intensity or parvalbumin-positive neuron number in the substantia nigra were analyzed using Cell Profiler image analysis software. Bars represent mean ± SEM, *n* = 4 mice per group. No significant differences between groups are detected by one-way ANOVA.

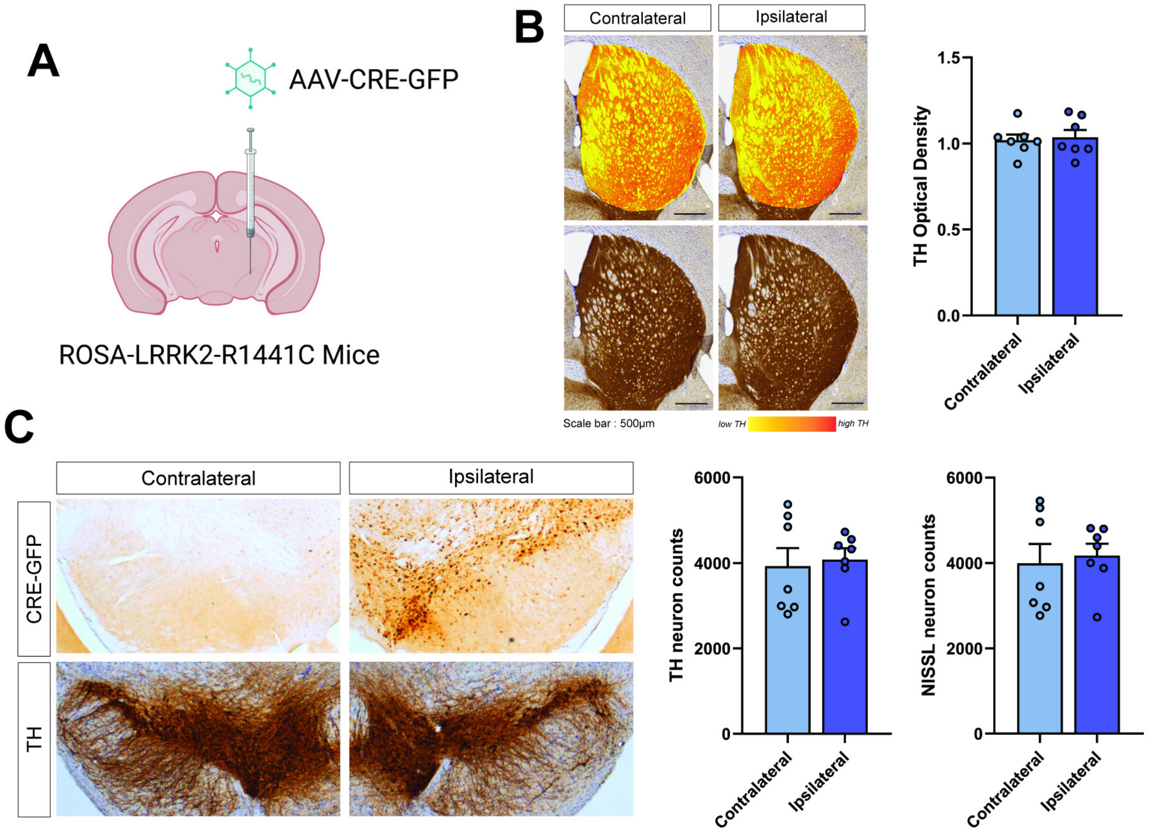


**Figure S3. Cre-GFP expression in the substantia nigra does not independently induce dopaminergic neurodegeneration in mice over 12 months.** (**A**) Homozygous floxed ROSA26-LRRK2-R1441C mice were unilaterally injected with AAV-Cre-GFP vector into the substantia nigra and analyzed after 12 months. (**B**) Immunolabeled images of TH-positive nerve terminals in the striatum, with optical density measured in the ipsilateral versus contralateral striatum using HALO analysis software. Bars represent mean ± SEM, *n* = 7 mice per group. (**C**) Immunolabeled images of TH-positive dopaminergic neurons or Cre-GFP-positive cells in the substantia nigra, with neurons counted by unbiased stereological analysis of TH-positive and total Nissl-positive neurons in ipsilateral versus contralateral substantia nigra. Bars represent mean ± SEM, *n* = 7 mice per group.

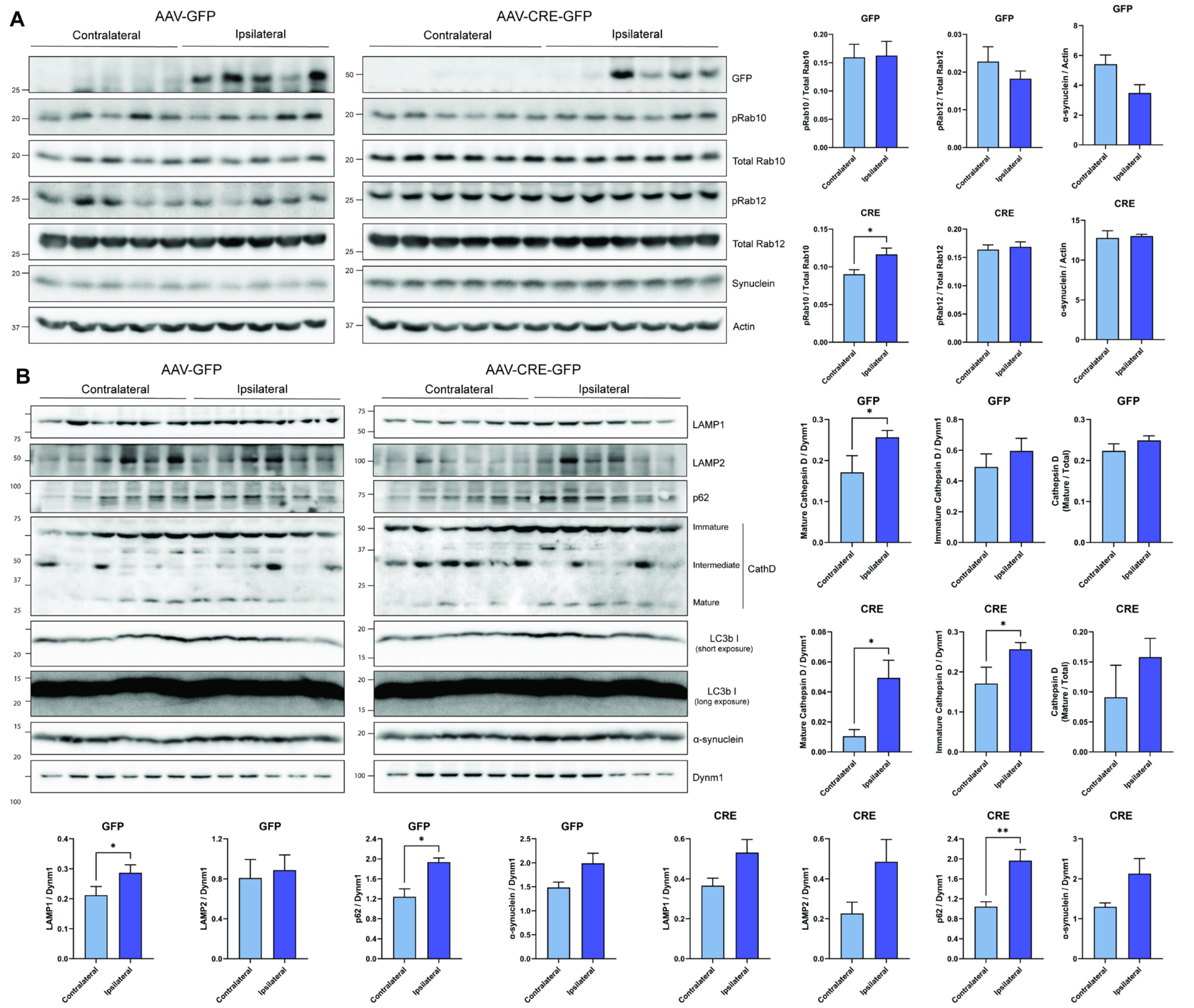


**Figure S4. Western blot analysis of autophagy-lysosomal pathway proteins in ventral midbrain tissue of *ATP13A2* KO mice at 3 months.** (**A**) 1% Triton-X100-soluble ventral midbrain extracts were prepared from *ATP13A2* floxed KO mice injected with AAV-GFP or AAV-Cre-GFP at 3 months. Western blots were probed for autophagy-lysosomal proteins, including pT73-Rab10, total Rab10, pS106-Rab12, total Rab12, total α-synuclein and actin as a loading control. Protein intensities were measured using ImageJ analysis software comparing ipsilateral and contralateral hemispheres for GFP or GFP-Cre mice. Bars represent mean ± SEM, *n* = 5-6 mice per group. $^{*}P<0.05$ by unpaired, Student's *t*-test. (**B**) 1% Triton-X100-insoluble (RIPA-soluble) ventral midbrain from the same mice. Western blots were probed for autophagy-lysosomal proteins, including LAMP1, LAMP2, p62, cathepsin D species, LC3B I and II, total α-synuclein and dynamin-

1 as a loading control. Bars represent mean ± SEM, *n* = 6 mice per group. **P*<0.05 or ***P*<0.01 by unpaired, Student's *t*-test.

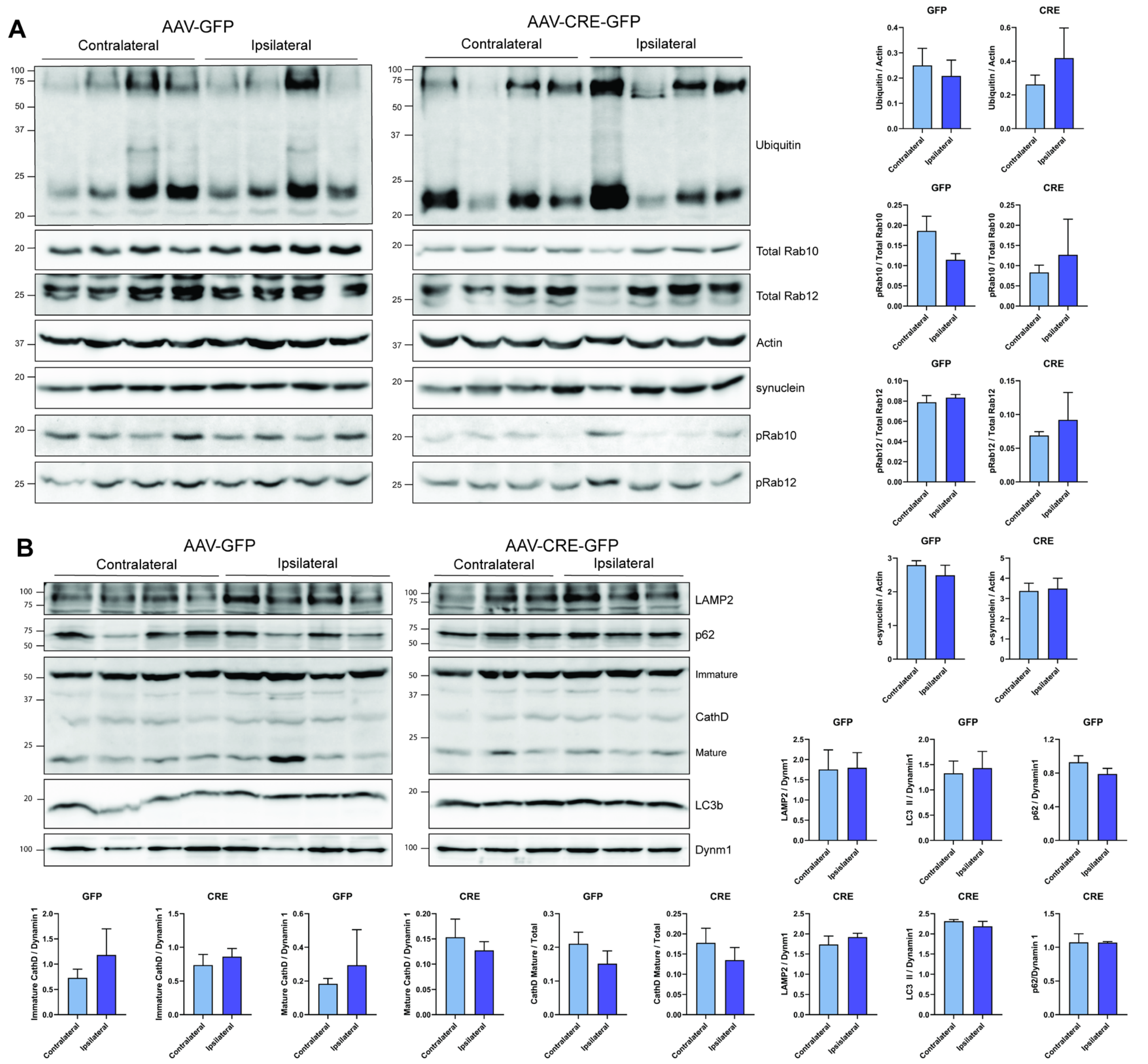


**Figure S5. Western blot analysis of autophagy-lysosomal pathway proteins in ventral midbrain tissue of *ATP13A2* KO mice at 6 months.** (**A**) 1% Triton-X100-soluble ventral midbrain extracts were prepared from *ATP13A2* floxed KO mice injected with AAV-GFP or AAV-Cre-GFP at 6 months. Western blots were probed for autophagy-lysosomal proteins, including total ubiquitin, pT73-Rab10, total Rab10, pS106-Rab12, total Rab12, total α-synuclein and actin as a loading control. Protein intensities were measured using

ImageJ analysis software comparing ipsilateral and contralateral hemispheres for GFP or GFP-Cre mice. Bars represent mean ± SEM, *n* = 4 mice per group. (**B**) 1% Triton-X100-insoluble (RIPA-soluble) ventral midbrain from the same mice. Western blots were probed for autophagy-lysosomal proteins, including LAMP2, p62, cathepsin D species, LC3B I and II, and dynamin-1 as a loading control. Bars represent mean ± SEM, *n* = 3-4 mice per group. No significance by unpaired, Student's *t*-test.
